# Knockdown of a novel ATPase domain of capsid protein inhibits genome packaging in potato leaf roll virus

**DOI:** 10.1101/2021.05.24.445413

**Authors:** Jitesh Kumar, Ravi Ranjan Kumar, Dilip Kumar Das, Auroshikha Mohanty, Kumari Rajani, Namaste Kumari, Vinod Kumar, Sunil Kumar, Tushar Ranjan

## Abstract

Potato leaf roll virus (PLRV) uses powerful molecular machines to package its genome into a viral capsid employing ATP as fuel. Although, recent bioinformatics and structural studies have revealed detailed mechanism of DNA packaging, little is known about the mechanochemistry of genome packaging in small plant viruses such as PLRV. We have identified a novel P-loop-containing ATPase domain with two Walker A-like motifs, two arginine fingers, and two sensor motifs distributed throughout the polypeptide chain of PLRV capsid protein (CP). The composition and arrangement of the ATP binding and hydrolysis domain of PLRV CP is unique and rarely reported. The discovery of the system sheds new light on the mechanism of viral genome packaging, regulation of viral assembly process, and evolution of plant viruses. Here, we used the RNAi approach to suppress CP gene expression, which in turn prevented PLRV genome packaging and assembly in *Solanum tuberosum* cv. Khufri Ashoka. Potato plants agroinfiltrated with siRNA constructs against the ATPase domain of CP exhibited no rolling symptoms upon PLRV infection, indicating that the silencing of CP gene expression is an efficient method for generating PLRV-resistant potato plants. Moreover, our findings provide a robust approach to generate PLRV-resistant potato plants, which can be further extended to other species. Finally, we propose a new mechanism of genome packaging and assembly in plant viruses.

## Introduction

Plant viruses cause diverse diseases in crops and are responsible for huge economic losses. The potato crop is severely affected by various biotic stresses. Among these stresses, viruses have a significant contribution to yield losses worldwide [1]. Due to vegetative reproduction of potato, viruses propagate through seed tubers and maintain their life cycle [2]. Tubers used for planting in the next season can harbor latent viruses, which subsequently reduce emergence, plant vigor, and yield [2]. More than 40 different viruses affect the cultivation of potato crop across the globe [1–2]. Potato leaf roll virus (PLRV), which belongs to the genus *Polerovirus* and family *Luteoviridae*, is a ubiquitous potato virus worldwide and is responsible for more than 20 million tons yield loss every year [3–4]. Thus, understanding the mode of infection, genome packaging, and assembly of PLRV is crucial for developing control strategies [4–5].

Genome packaging is an important step in the process of viral maturation. During virus life cycle, genetic information needs to be incorporated into newly produced virus particles [6–7]. Three types of genome packaging system have been reported in viruses so far [5–6]. In the passive type I system, which exists in majority of plant viruses, the capsid proteins (CPs) nucleate over the genome and form a mature virus particle without utilizing any energy. On the contrary, both type II and III systems, which are found in bacteriophages and large eukaryotic DNA viruses, are active, ATP-dependent packaging systems [5–7]. Viruses from type II and III system use powerful ATPase motors to achieve genome packaging [6]. These motors generate forces as high as 100 pN and translocate DNA into a preformed prohead until DNA condenses into crystalline form [8]. These viral packaging motors share a common architecture with P-loop superfamily of multimeric ring ATPases that perform diverse functions such as chromosome segregation (helicases), protein remodeling (chaperones), and cargo transport (dyneins) [9].

Recently, we proposed a novel, expanded sub-classification system for the type I genome packaging system and placed many of the plant viruses, including PLRV, under the ATP-dependent sub-type IA system based on the ATPase domain present in CP [5]. The mechanochemistry of the ATPase motor encoded by plant viruses is hardly explored. How a functional CP ATPase oligomerizes and nucleates on the genomic end and initiates its activity is not yet known. The ATPase motors drive the difficult task of precisely inserting the viral genome into the capsid from the cytoplasmic pool of RNA inside the host cell. Presence of a powerful motor across viruses infecting different domains of life might represent a unique variation of the genome packaging apparatus acquired and contrived by plant viruses [7].

As CP is known to play a crucial role in genome packaging in PLRV [5], we sought to analyze its amino acid sequence to identify conserved motifs and use them to develop a potential method of developing virus-resistant plants. Our comprehensive bioinformatic analysis of CP amino acid sequence revealed a unique P-loop containing ATPase domain with two Walker A-like motifs, two arginine fingers, two sensor-like motifs, and one Walker B motif, which possess ATP-binding and hydrolysis activities and are spread throughout the polypeptide chain of CP. Our structural analysis revealed that these critical motifs are either part of the loop or present at the tip of strand and make a flexible active site for ATP binding and hydrolysis. The P-loop plays a key role in coordinating ATP hydrolysis with DNA translocation. We believe that the ATPase domain of CP has a direct role in genome packaging of PLRV. To validate this hypothesis, we conducted RNAi experiments on the CP gene to check its effects on PLRV infection. Knockdown of the ATPase domain of the CP gene resulted in generation of potato leaves free from PLRV, indicating its role in genome packaging and assembly. Further, northern blotting, RT-PCR, and ELISA analyses confirmed that the PLRV CP mRNA was not detected in the leaves of plants agroinfiltrated with CP siRNA. However, the same mRNA was detected at high levels in the tertiary leaves of control plants (naturally infected) and those agroinfiltrated with the empty vector. As CP homologs are present in almost every plant virus, their inhibition via RNAi can be a potential method for generating virus-resistant plants.

## Experimental procedures

### Sequence analysis, motif identification and structural analysis

The sequences of CP from different strains of PLRV were retrieved from the NCBI database (http://www.ncbi.nlm.nih.gov/). Multiple sequence alignments were generated using ClustalW [10] and were manually corrected for domain superimpositions. Atomic structure and secondary structure of CP were predicted using Iterative Threading ASSEmbly Refinement (I-TASSER), a tool which builds atomic structures based on iterative template fragment assembly simulations and multiple-threading alignment using Locally Installed Meta-Threading Server (LOMETS) [11]. This prediction returns structures with good C-scores in the range of −2.04 to −1.09.

### Preparation of siRNA constructs

To generate target-specific siRNAs against the CP gene, the CP-encoding gene fragments were amplified using the cDNA of PLRV as template. The amplified fragments were digested with *Xho I* and *Kpn I* and cloned into the pHANNIBAL vector at *Xho I* – *Kpn I* sites in sense orientation (pHANNIBAL-sense CP). The primer sequence for cloning of the CP-encoding gene fragment in sense orientation was designed manually and synthesized at IDT, USA. These sequences are as follows: CPs FP: 5’-CCGG*<u>CTCGAG</u>*ATGAGTACGGTCGTGGTTAGAGG A-3’; CPs RP: 5’-GCGC*<u>GGTACC</u>*CTATCTGGGGTTCTGCAAAGCCAC-3’. The pHANNIBAL vector contains the CaMV 35S promoter and the NOS terminator in sense orientation. Subsequently, the amplified and digested antisense CP gene fragments were cloned into the same vector at *HindIII*– *Xba I* sites to generate the antisense construct (pHANNIBAL-antisense CP). The sequences of these primers are as follows: CPas FP: 5’-GGGT*<u>AAGCTT</u>*CTATCTGGGGTTCTGCAAAGCCAC-3’; CPas RP: 5’-GCGC*<u>TCTAGA</u>*ATGAGTACGGTCGTGGTTAGAGG-3’. The resulting siRNA construct containing the sense and antisense fragments of the CP cDNA sequence was named as pHANNIBAL-CP. The siRNA cassettes were released from pHANNIBAL-CP by *Not I* digestion and introduced into the binary vector pART27 to generate the siRNA construct pART27-CP for plant transformation. Subsequently, the orientation of the cloned CP genes was confirmed through sequencing at each step.

### Transient expression assay

The binary plasmid pART27-CP and the empty vector pART27 were extracted and purified from *E. coli* cultures and were transformed into *Agrobacterium tumefaciens* EHA101 using the freeze–thaw transformation method [12]. The transformed cells of *A. tumefaciens* were plated on Luria-Bertani (LB) agar plates containing 50 μg/mL kanamycin, 100 μg/mL spectinomycin, and 100 μg/mL chloramphenicol for selecting successful transformants. Transformation of the binary plasmid was further confirmed using colony PCR for the CP gene. For agroinfiltration, *A. tumefaciens* cells harboring the pART27-CP siRNA constructs were grown overnight at 28°C in the LB medium supplemented with appropriate antibiotics. The overnight cultures were diluted 1:10 in fresh media containing the above-mentioned antibiotics, 10 mM 2-(N-morpholino) ethanesulfonic acid (MES), and 200 μM acetosyringone to reach an OD_600_ of 0.3. The cells were collected by centrifugation at 5000 ×*g* for 5 min and were resuspended in the infiltration medium containing 10 mM MES, 10 mM MgCl_2_, and 200 μM acetosyringone. The cells were then incubated at room temperature for 2–3 h before agroinfiltration. *Solanum tuberosum* cv. Khufri Ashoka plants were agroinfiltrated by injecting 2 mL of these cells directly into the phloem and leaves using a syringe. Whole plants were covered with a transparent plastic bag for 3–4 days. All experiments were repeated thrice with five plants in each experiment for each siRNA construct and control.

### Viral infectivity assay

Symptomatic potato plants newly emerged from PLRV-infected tubers were used for agroinfiltration of pART27-CP siRNA constructs. PLRV-infected tubers were grown in a controlled environment of transgenic glass room. The presence of PLRV titer in tubers and newly emerged plants were confirmed through enzyme-linked immunosorbent assay (ELISA) and PCR using CP-specific primers (FP: 5’ATGAGTACGGTCGTGGTTAGA-3’; RP: 5’ CTATCTGGGGTTCTGCAAAGCC-3’). The plants were grown in a transgenic glass house at 25 °C with 16 h of light and 8 h of darkness after agroinfiltration. Ten days post agroinfiltration (dpi), newly emerged tertiary leaves were harvested for RNAi analysis. The RNAi constructs were analyzed using Northern blotting, RT-PCR and ELISA.

### Northern blotting analysis of siRNA

Leaves of 5–10 dpi agroinfiltrated potato plants and healthy control plants were used for small RNA isolation. Small RNA was extracted using PAX gene Tissue RNA/miRNA Kit (Qiagen, Germany) following the manufacturer’s protocol. Denaturing polyacrylamide gels (15%) containing 7 M urea were run at 40 mA (600 V) for approximately 2 h to resolve the total RNA, which contained the siRNA pool. The gels were stained for 10 minutes with 0.5 μg/ml ethidium bromide (EtBr) in DEPC-treated TBE buffer. A Bio-Rad transblot apparatus (Bio-Rad, USA) was used to transfer the RNA onto a Hybond-N + positively charged nylon membrane (GE healthcare, USA) at 200 mA (9–10 V) for 3 h followed by crosslinking for 20 min at 1200 μJ and drying for 30 min at 50 °C to improve sensitivity. Further, the membrane was prehybridized in pre-hybridization buffer (7 % SDS, 200 mM Na_2_HPO_4_ (pH 7.0), 5 μg/ml salmon sperm DNA (SSDNA)) at 40 °C for 30 min. The hybridization buffer was replaced with hybridization buffer containing biotin-labeled probes at 50 pmol/mL concentration. The membrane was subsequently hybridized at 40 °C for 12 to 16 h with continuous gentle shaking followed by rinsing with washing buffer (1× SSC, 0.1% SDS) thrice for a total of 15 minutes at room temperature. The membrane was then blocked for 15 minutes with blocking buffer using gentle shaking at room temperature followed by incubation with hybridization buffer containing stabilized streptavidin-HRP conjugate for an additional 15 minutes. The blot was visualized using Amersham typhoon blot imaging systems (GE, Healthcare) after developing by ECL (GE Healthcare).

### RT-PCR analysis of viral RNA in potato plants

Total RNA was extracted from the newly emerged tertiary leaves (100 mg) of agroinfiltrated potato plants and healthy control plants using RNeasy Plant Mini kit (Thermo Scientific, USA) following manufacturer’s instructions. RT-PCR was conducted for CP-infiltrated plants individually using the PLRV-specific primers. cDNA was synthesized using an RT-PCR kit (Thermo Scientific, USA).

### ELISA

Indirect ELISA was performed for the detection of PLRV infection. Tertiary leaves of agroinfiltrated plants and healthy control plants (positive and negative) (0.5 g) were macerated in 5 mL extraction buffer (0.05 M phosphate-buffered saline, 0.01 M sodium diethylene carbamide, pH 7.4). As positive and negative controls, PLRV infected and uninfected/healthy potato leaves were considered, respectively. The extracts were centrifuged at 10,000×*g* for 5 min. One hundred microliter of the supernatant was added to the ELISA plate, which was incubated overnight at 4 °C. Further, 200 μL of rabbit polyclonal antibody against PLRV-CP (1:5000; Promega) was added to each well and incubated for 3 h at 37 °C. After washing, goat anti-rabbit immunoglobulin G conjugated with alkaline phosphatase (Promega, USA) was added to the wells and incubated for 3 h at 37 °C. The substrate p-nitrophenyl phosphate was added to the plate for developing, and absorbance at 405 nm was measured. The absorbances of samples and controls were normalized to that of the positive control.

## Results

### Insights on the structural details of the PLRV CP ATPase domain

CPs are known as plant virus-encoded factors that nucleate over viral genomes and encapsidate them by deriving energy from ATP [5]. The motifs necessary for ATP binding and hydrolysis are present throughout the polypeptide chain of PLRV CP (Fig 1A & B). Fig 1A presents a multiple sequence alignment of CP sequences from different PLRV variants. Intriguingly, our sequence analysis revealed a unique pattern of ATPase domain, which contained two sets of Walker A, arginine finger, and sensor-like motifs, suggesting divergent evolution within the classical P-loop-containing ATPase superfamily. The Walker-A “P-loop” motif is proposed to coordinate ATP hydrolysis with DNA translocation. The Walker A-like motif (lavender color), with the consensus sequences RGRGSSET, interacts with the β- and γ-phosphates of the bound ATP, whereas the conserved Asp of Walker B (consensus sequence hhhhDG), located next to a β-strand, binds to a metal ion and helps in ATP hydrolysis. The Walker-B (blue) Asp coordinates with the Mg^2+^ cation, while another conserved catalytic Gly residue primes a water molecule for the nucleophilic attack on the γ-phosphate of the bound ATP. Our sequence and structure analysis found that all the critical motifs are either present at the tip of the strand or part of the loop, indicating relatedness of CPs to ASCE P-loop ATPases (Fig 1A). Further, we observed that the arginine fingers I and II (red), which are located 9-10 residues downstream of sensor I (green) motif and 5-6 residues downstream of Walker A’ (lavender), and the sensor motif II (green) present about five residues further downstream of the Arginine finger II motif are strictly conserved across various strains of PLRV analyzed in this study (Fig 1A). Sensor motif I (green) is located 48 residues downstream of Walker B and flanking of arginine finger I, II and Walker A’ with sensor motifs I and II represents the uniqueness of this novel P-loop-containing ATPase. Our structure prediction using I-TASSER revealed that the active site of PLRV CP comprises all the motifs necessary for ATP interaction except Arginine finger I, which is situated far from the rest of the motifs in their folded structure (highlighted in red; Fig 1B). Arginine finger is a classical hallmark of ATPases and is conserved across many ATPases. It completes the active site from a distinct location, forming contacts with the γ-phosphate of the nucleotide [13].

**Fig 1.**
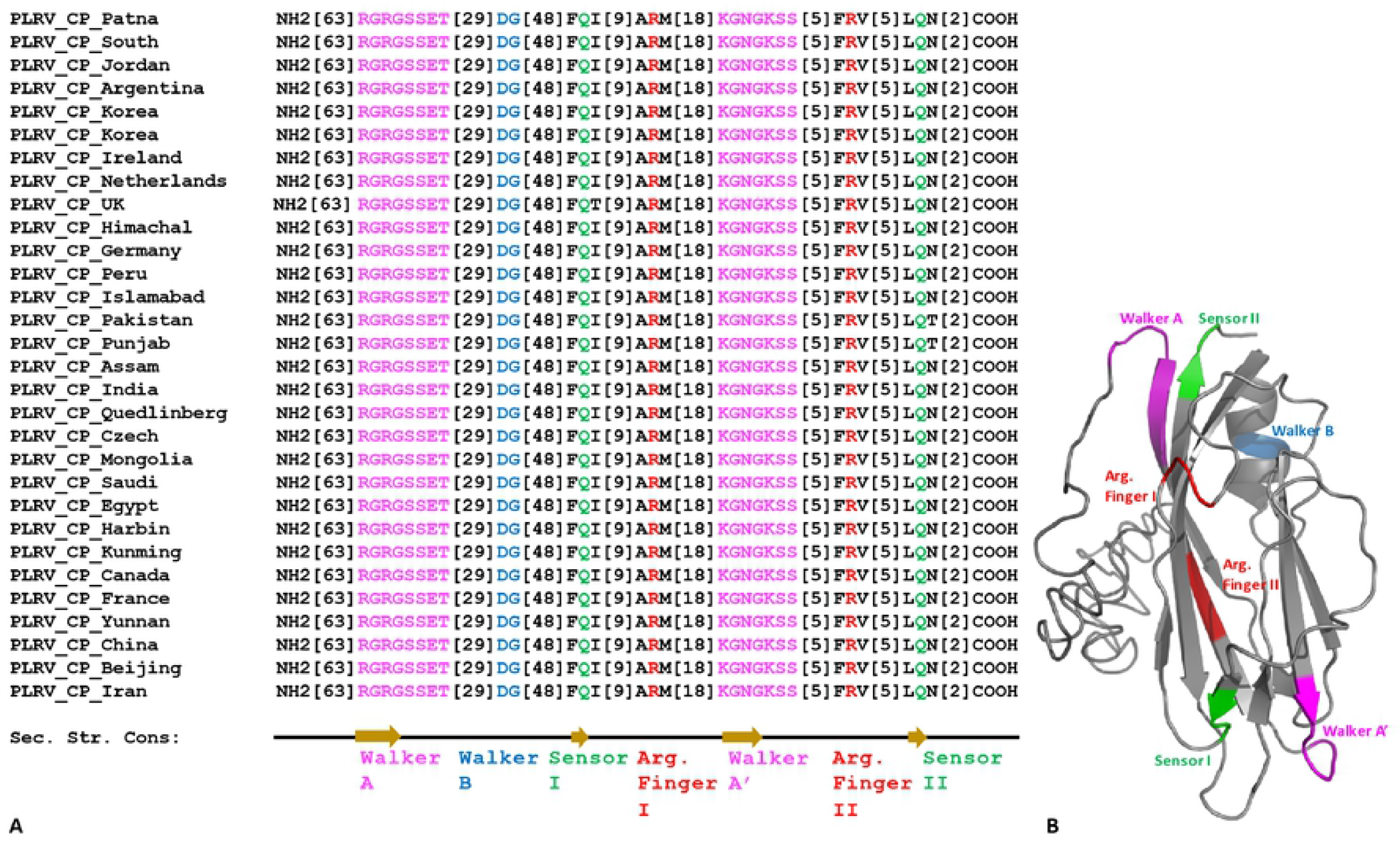
(A) Multiple sequence alignment of the ATPase domain of CP from different variants pf PLRV. The alignment was generated using Clustal W and manually corrected for domain superimpositions. The number(s) in brackets represent the number of amino acids. (B) I-TASSER predicted atomic model of PLRV CP with all the ATP catalysis motifs (Walker A and A’ in lavender; Walker B in blue; Sensor I & II in green and arginine finger I & II in red color). [Accession Numbers: PLRV China: QEE13716.1, PLRV Shimla: AFJ11896.1, PLRV Beijing: QEE13717.1, PLRV Japan: BBN91709.1, PLRV Yunnan: ACO92389.1, PLRV France: AAL77948.1, PLRV Saudi Arabia: AGR87640.1, PLRV Iran: ACK99523.1, PLRV Mongolia: AHA43773.1, PLRV Pakistan: ASB17109.1, PLRV Punjab: ASB17109.1, PLRV Canada: AYA73305.1, PLRV Netherlands: CAA54535.1, PLRV Czech Republic: ABY49848.1, PLRV Kunming: ACO92404.1, PLRV Harbin: AKN09918.1, PLRV UK: CAA54537.1, PLRV India: AFJ11862.1, PLRV Germany: QBO24571.1, PLRV Himachal Pradesh: AAN31764.1, PLRV Islamabad: ASB17111.1, PLRV Jordan: ACF33055.1, PLRV Quedlinberg: AFI93516.1, PLRV Ireland: QJF11910.1, PLRV Korea: AAG14887.1, PLRV South Africa: AAB80766.1, PLRV Korea: AAD00221.1, PLRV Argentina: ARS33719.1, PLRV Egypt: ATV90882.1, PLRV Assam: QDA76478.1, PLRV Patna: MW027216.1, PLRV Peru: APC60287.1]

### Construction of the pART27 binary vector

Specific primers were designed for the amplification of the gene sequence of CP in sense and antisense orientations. The generation of siRNA construct is illustrated in Fig 2A. The total RNA extracted from infected potato leaves was subjected to RT-PCR to generate cDNA, and the desired sequences were amplified from the cDNA using oligo dT primers. This PCR yielded two 627 bp bands: one of sense orientation and another antisense (Fig 2B; lanes 1 and 2). These fragments were cloned into the pHANNIBAL vector (5.8 kb) after digestion with the appropriate set of enzymes (Fig 2C; lane 1). The cloning of the CP gene in pHANNIBAL in the sense orientation (pHANNIBAL-sense CP) and antisense orientation (pHANNIBAL-antisense CP) was carefully analyzed using different sets of restriction enzymes (Fig. 2D; lanes 1-7). Further, the orientation of genes was confirmed through sequencing (IDT). Another round of sequencing was performed for the cloning vector carrying both sense and antisense CP (pHANNIBAL-CP). The resulting siRNA constructs were further digested by *Not I* and introduced into the digested linear (~11.6 kb) binary vector pART27 (Fig 2E; lane 1) to generate the pART27-CP-siRNA construct. The accuracy of siRNA constructs in pHANNIBAL (Fig 2D) and pART27 (Fig 2F) binary vector was further confirmed through restriction digestion analysis and sequencing.

**Fig 2.**
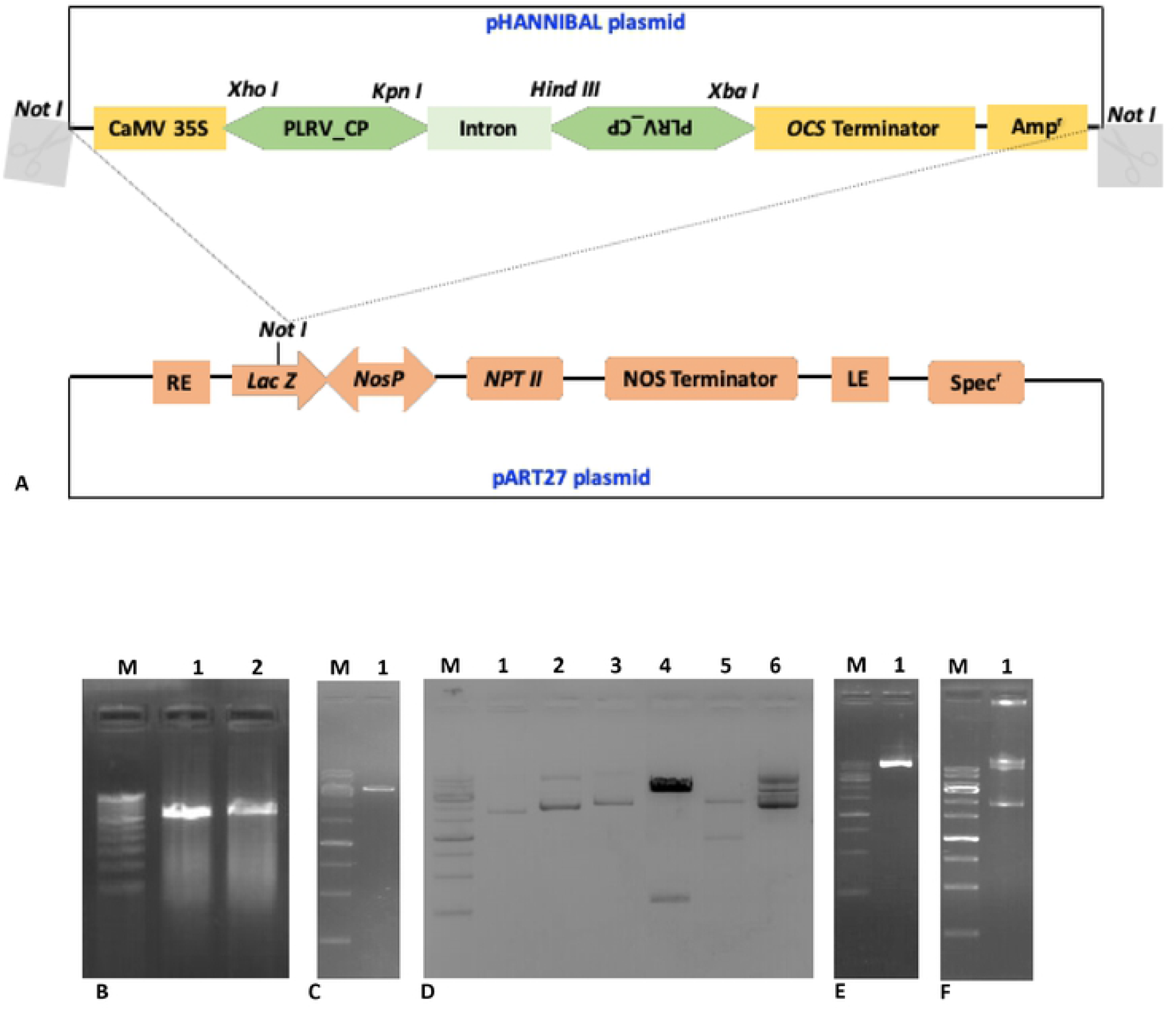
(**A**) Schematic representation of the generation of pART27-CP siRNA constructs used for transient expression by agroinfiltration. CP genes were first cloned in pHANNIBAL plasmid in sense and antisense orientations. Subsequently, the complete gene cassette was transferred into the binary pART27 plasmid. **(B)** RT-PCR was performed to amplify sense and antisense PLRV CP sequences from the leaves of infected potato plants. Amplified cDNA fragments were analyzed by electrophoresis on 0.8% agarose gel. M: 100 bp DNA ladder; lanes 1 and 2: PCR products of sense and antisense CP (627 bp). (**C**) The pHANNIBAL plasmids were purified from *E. coli* DH5α cultures and analyzed by electrophoresis on 0.8% agarose gel. M: 1 kb DNA ladder; lanes 1: pHANNIBAL plasmid (5.8 kb). (**D**) Confirmation of siRNA constructs in pHANNIBAL and pART27 binary vector through restriction analysis. M: 1 kb DNA ladder; lane 1: undigested pHANNIBAL; lane 2: antisense CP ligated with pHANNIBAL (pHANNIBAL-antisense CP construct); lane 3: both antisense and sense CP ligated with pHANNIBAL (pHANNIBAL-CP-siRNA construct); lane 4: *Hind III* and *Xba I* digestion of pHANNIBAL-antisense CP construct shows the release of a 627 bp fragment; lane 5: *Xho I* and *Xba I* digestion of pHANNIBAL-CP siRNA construct shows the release of a~ 1500 bp fragment; and lane 6: *Not I* digestion of pHANNIBAL-CP siRNA construct shows the release of two fragments of ~ 4 kb (including sense CP, intron, and antisense CP sequence) and 3.5 kb. (**E**) The pART27 binary plasmid was purified from *E. coli* DH5α culture and analyzed by electrophoresis on 0.8% agarose gel. M: 1 kb DNA ladder; lane 1: pART27 plasmid (11.6 kb). (**F**) The whole siRNA cassette (~ 4 kb) was transferred from pHANNIBAL to pART27, and this was further confirmed by *Not I* restriction analysis of recombinant pART27.

### Transient expression of CP siRNA

To determine whether the knockdown of CP could interfere the genome packaging and assembly of PLRV in potato, the pART27-CP-siRNA constructs were agroinfiltrated into *S. tuberosum* cv. Khufri Ashoka. Control plants (PLRV infected) (Fig 3A; lane 1) and those agroinfiltrated with the empty vector (Fig 3A; lane 2) showed rolling symptoms of PLRV infection in the upper leaves, whereas plants that were agroinfiltrated with the CP siRNA constructs (pART27-CP) did not show any symptoms of viral infection (Fig 3A; lane 5). Rolling symptoms were also observed in the upper leaves of plants agroinfiltrated with pART27-sense CP and pART27-antisense CP construct (Fig 3A; lane 3 & 4 respectively). These findings indicated that siRNA constructs suppressed viral genome packaging and assembly in the plants.

**Fig 3.**
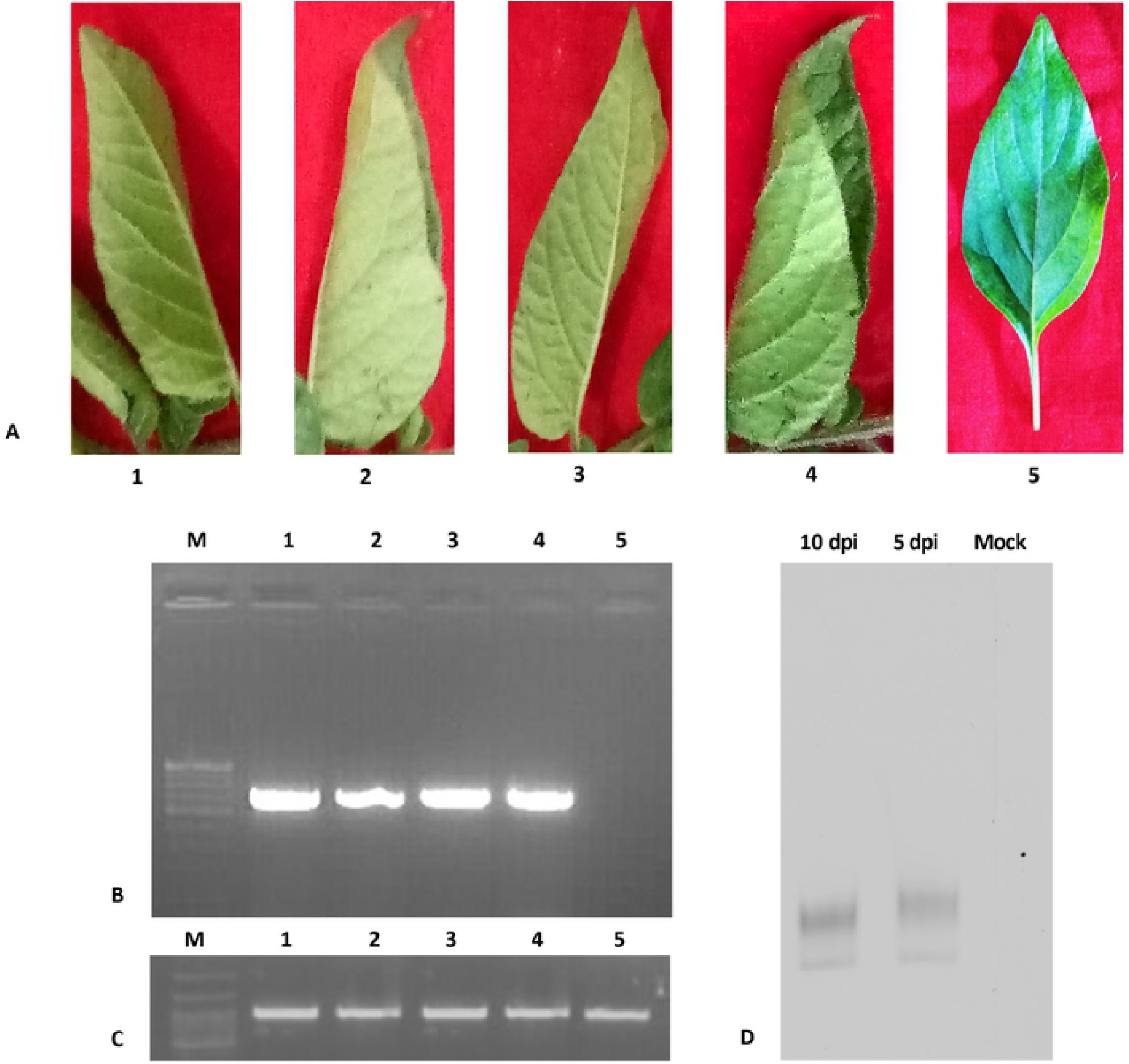
**(A)** Symptoms observed in the tertiary leaves of the PLRV infected potato plants. (1) PLRV-infected control without agroinfiltration; (2) agroinfiltrated with the empty vector pART27; (3) agroinfiltrated with the plasmid containing only antisense sequence (pART27-antisense CP); (4) agroinfiltrated with the plasmid containing only sense sequence (pART27-sense CP); (5) agroinfiltrated with the pART27-CP siRNA construct; **(B)** Detection of PLRV RNA by RT-PCR. Amplified cDNA fragments (627 bp) were analyzed by electrophoresis on 0.8 % agarose gel. M: 100 bp marker; lane 1: PLRV was observed in control plant; lane 2: PLRV from tertiary leaves of the plant containing the empty vector pART27; lane 3: PLRV from tertiary leaves of the plant containing the antisense construct (pART27-antisense CP); lane 4: PLRV from tertiary leaves of the plant containing the sense construct (pART27-sense CP); lane 5: no PLRV in tertiary leaves of the plant containing CP siRNA (pART27-CP). **(C)** Actin PCR was performed as an internal control. **(D)** Confirmation of expression of siRNA using Northern blotting analysis. Higher expression of siRNA in the leaves at 10 dpi (lane 1) was observed as compared to 5 dpi (lane 2), whereas empty vector (mock) agro-infected leaves did not show any siRNA (lane 3).

### RT-PCR analysis of the CP gene

To detect the presence of the virus in the host tissue, total RNA isolated from the upper tertiary leaves of agroinfiltrated plants and control plants was subjected to RT-PCR using PLRV-specific primers. The 627 bp amplified DNA fragment corresponding to the CP gene of PLRV was detected in the tertiary leaves of plants agroinfiltrated with the empty vector and control leaves; however, it was not detected in the tertiary leaves of plants agroinfiltrated with the CP siRNA constructs. Thus, RT-PCR confirmed that PLRV infection was suppressed by these siRNA constructs (Fig 3B). RT-PCR for the actin gene was also performed simultaneously as an internal control (Fig 3C).

### Analysis of agroinfiltrated potato plants harboring siRNA constructs

The expression of siRNA was analyzed by Northern blotting using an equal amount of total RNA from the leaf samples of pART27-CP construct and empty vector harboring plants respectively, to ascertain whether RNA silencing was initiated in agro-infected potato leaves by pART27-CP construct. The agro-infected regions of the leaf were used to isolate total RNA, which was subsequently analyzed for the accumulation of siRNAs at 5 and 10 dpi. The siRNAs were detected in the agro-infected region at 5 and 10 dpi (Fig 3D). The RNA band intensity suggested higher expression of siRNA in the leaves at 10 dpi (Fig 3D; lane 1) than that at 5 dpi (Fig 3D; lane 2), whereas empty vector (mock) agro-infected leaves did not show any presence of siRNA (Fig 3D; lane 3).

### ELISA for detecting PLRV in host plants

The presence of PLRV in host plants was detected by DAS-ELISA using an anti-PLRV CP antibody. The absorbance values of all the samples were normalized to that of the positive control. The mean absorbance values for all the samples are presented in Fig 4A. We noted that the relative absorbance values of the samples from plants containing the CP siRNA construct were similar those of the negative controls (healthy plants). Overall, PLRV CP was undetectable in plants agroinfiltrated with siRNA, whereas PLRV was present in high concentration in the plants agroinfiltrated with the empty vector, with only the sense construct, or only the antisense construct.

**Fig 4.**
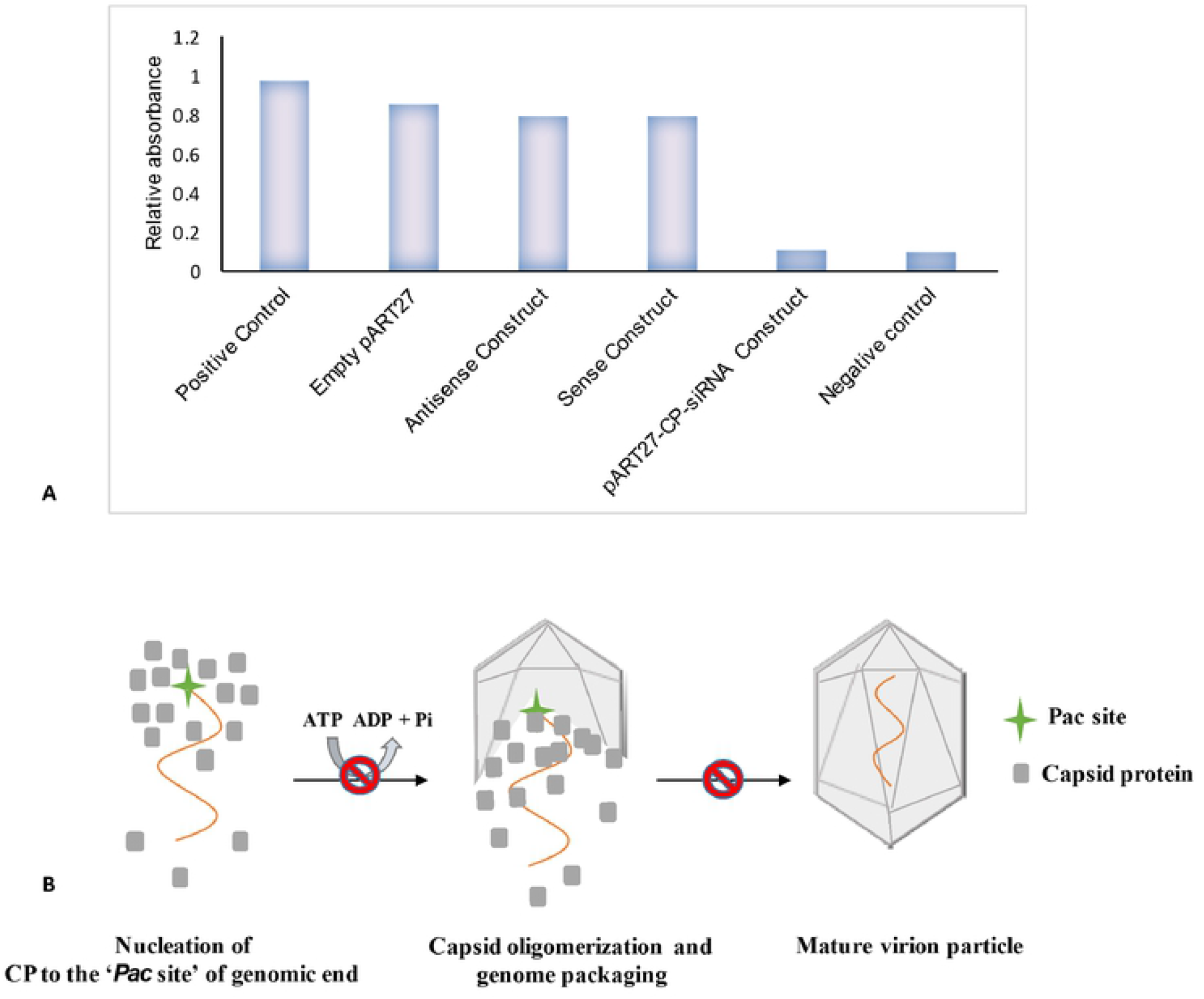
**(A)** ELISA to determine the titer of PLRV. Absorbance values of all the samples were normalized to that of the positive control (PLRV infected plant). A healthy uninfected plants were considered as negative control. **(B)** Hypothetical model for the genome packaging and assembly in PLRV. CP recognizes the end of the genome (*pac site*) and nucleates over the genome. CP further oligomerizes and encapsidates the unit length genome and leads to the formation of mature virion particle. Knockdown of the ATPase domain of CP resulted in either inhibition of genome packaging or generation of non-functional/immature virus particle.

## Discussion

Mechanism of genome packaging is reasonably well understood in viruses belonging to the type II and III packaging systems [6–7, 14–16]. However, the detailed mechanism of genome packaging in plant viruses with type I packaging system remains unclear. Whether plant viruses should be placed in a group of energy-dependent packaging system or energy-independent one was debatable until we proposed a subgroup of the type I system. In this subgroup, which we named as type IA, majority of plant viruses such as PLRV, potato virus X, and other geminiviruses were finally classified [5–6]. The plant viruses that possess ATPase domain in their CP were placed under this subgroup and for these viruses, direct role of CP in genome packaging was hypothesized [5]. Our sequence analysis (Fig. 1A) and atomic structure prediction (Fig. 1B) of CP revealed the presence of a novel P-loop-containing ATPase motif. The Walker-A motif is a phosphate-binding loop (P-loop) found in possibly the most ancient and abundant protein class, so-called P-loop ATPases. Researchers have proposed that the Walker-A “P-loop” motif coordinates ATP binding and hydrolysis with DNA translocation during active genome packaging in plant viruses and that it is a part of the ATPase motor [18–19]. During the assembly of viruses, a powerful ATP-driven motor translocates and packages DNA into a capsid coat. The crucial Arginine finger I motif is distantly placed in the atomic model from the two Walker A, B, sensor, and arginine finger II motifs, which form a cleft or active site for the binding of ATP. Once ATP occupies its position in the active site, the arginine finger comes into the vicinity of the γ-phosphate for hydrolysis. The unique arrangement of ATP-binding and hydrolysis motifs in the primary and tertiary structure of CP (Fig. 1A & B) indicates the structural similarity of CP with the members of the well-known classical P-loop ATPase superfamily [17–19]. The role of two Walker A-like motifs (i.e., Walker A and Walker A’) during ATP binding and catalysis needs to be experimentally investigated. Replacing the critical amino acid residues of these ATPase motifs through site directed mutagenesis (SDM) is underway to further dissect the function of these motifs, especially in terms of active genome packaging in assembly.

In the present study, the efficiency of CP siRNA constructs to silence and inhibit PLRV assembly was assessed. The siRNA constructs were designed against CP of PLRV and agroinfiltrated into infected potato plants. The agroinfiltrated plants did not show PLRV infection. Suppression of viral infection could be attributed to the reduced expression of CP due to its silencing by the siRNA constructs. The present study indicated that the transient expression of CP constructs resulted in specific and efficient inhibition of PLRV. Northern blotting, RT-PCR, and ELISA also confirmed that the PLRV CP mRNAs were not detected in the leaves of plants agroinfiltrated with CP siRNA, whereas these mRNAs were detected at high levels in tertiary leaves of control plants (naturally infected) and those agroinfiltrated with the empty vector. These findings further confirm the earlier reports and suggest that CP ATPase has a direct role in the genome packaging, and suppression of genome packaging may lead to genome deficient and non-functional viral particles [5,20]. Thus, the presence of three basic types of viral DNA packaging motors probably indicates independent innovations.

To the best of our knowledge, this is the first study that presents a novel P-loop containing ATPase fold of CP comprising several repeat motifs and demonstrates that the suppression of CP expression using siRNA constructs can lead to resistance against PLRV. The inhibition of the expression of CP by gene silencing is an efficient and promising method to introduce resistance to PLRV [2, 4–5, 21]. As PLRV causes severe crop yield losses in the potato growing regions worldwide [22–23], our findings will be helpful for developing PLRV-resistant potato crops. This approach can also be applied to an extensive range of plant species to develop resistance against various viral diseases. We are developing strategies for the development of PLRV-resistant potato varieties using the RNAi approach by targeting multiple genes.

## Acknowledgement

This work was supported by grants from the Science and Engineering Research Board (SERB), Department of Science and Technology, Govt. of India, New Delhi (India) (File No. SRG/2019/002223) to TR. We would also like to thank Bihar Agricultural University, Sabour, for providing the basic infrastructure for conducting the research works.

## Competing interests

The authors have declared no competing interests exist.

## Data Availability

Not applicable

